# Erythropoietic, hematinic and leucocytic activities of the aqueous extracts of the fruits and leaves of *Solanum torvum*

**DOI:** 10.1101/2023.04.04.535631

**Authors:** Ilyas Ibrahim, Denis Bardoe, Daniel Hayford

## Abstract

The Turkey berry *(Solanum torvum)* plant is globally exploited for its medicinal and pharmacological benefits. This study was conducted to investigate the erythropoietic, hemanitic and leucocytic activities of the aqueous extracts of the fruits and leaves of *Solanum torvum*. Three groups of rabbits with five replicates each (aged 6-8 weeks) where either given the aqueous extracts of fresh turkey berry fruits, fresh turkey berry leafs, or distilled water at a dose of 0.5ml and a concentration of 100g of leaf or fruit against 100ml of distilled water (1g per 1ml of distilled water) per day over a six-week period. The rabbits were fed *ad libitum* with concentrate and water throughout the experimental period. A full blood count (FBC) using a BC 2600 hematology analyzer was conducted on the collected blood samples of the experimental animals at the end of the sixth week of extract administration. The blood samples of the rabbits given the aqueous extracts of the fruit and leaf of turkey berry at the end of the six weeks of treatment administration showed a significant (p<0.05) increase in red blood cell count, hemoglobin level, white blood cell count and hematocrit. The results indicated that *Solanum torvum* had a positive erythropoietic, hemanitic and leucocytic properties. This implies that *Solanum torvum* is effective in improving the quality of blood and hence can prevent some hematological disorders.

## 1. Introduction

*Solanum torvum* commonly referred to as turkey berry is a wild plant, which is widely used mostly in Africa and Asia for its pharmacological and nutritional benefits. The plant remains one of the most mysterious plants in terms of the proven thoughts about it until today despite its wide geographical occurrence, usage and documented scientific facts (Ogah, 2015). Turkey berry has been for centuries used in the treatment of a number of diseases viz. anemia, ulcer, asthma, hypertension, colds, cough etc. (Yousaf *et al*., 2013). Very few of the numerous perceptions about turkey berry are documented until recently; most of the uses of the plant were based on myths and traditions with no backing proofs. In Ghana, turkey berry happens to be the most recommended recipe by doctors, midwives and other traditional healers for people who have anemia and other related cases (Ogah, 2015). Kuffuor *et al*., (2011) demonstrated the Immunomodulatory and erythropoietic effects of turkey berry plant on Sprague–Dawley rats when they induced anemia in the rats by phenyl hydrazine (PHZ). Chah *et al*., (2000) explained the antimicrobial and antifungal activities of the methanolic extract of *Solanum torvum* by treating it against a number of bacterial and fungal species isolated from the clinical samples of humans and animals by disc fusion. Akoto *et al*., (2015), accessed the mineral and nutritional compositions of *Solanum torvum* fruits from Ghana and found it to contain iron (76.869mg/kg), manganese (19.466mg/kg), calcium (221.583mg/kg), copper (2.642mg/kg) and zinc (21.460mg/kg). Additionally, Vitamins A and C contents were also analyzed and found to be 0.078mg/100g and 2.686mg/100g respectively. Anemia has negative effects on the health and economic wellbeing of nations and communities (Ghana National anemia profile, GNAP, 2014). Children with anemia experience irrevocable cognitive and developmental delays and exhibit decreased worker productivity as adults (Walker, *et. al*, 2007). Globally, maternal anemia increases the risk of pre-term delivery and low birth weight, and iron-deficiency anemia underlies 115,000 maternal deaths and 591,000 perinatal deaths each year (Stoltzfus, Mullany, and Black. 2004). According to the GNAP (2014), the percentages of women (Aged 15-49 years) and Children (Aged 5-65 months) with mild and moderate anemia were 42.4% and 65.7% respectively in 2014.. There is a need therefore to research in how iron-rich plants such as *S. torvum* can help undo the global disasters caused by anemia. Again, although some of the pharmacological and medicinal properties of *Solanum torvum* are documented. Among the least observed areas are its erythropoietic and hemanitic effects. Moreover, very few reports are having been made on the effects of *Solanum torvum* on other hematological properties such as on leucocytes. Therefore, there is a need for further explorations in this area.

## 2. Materials and Methodology

### 2.1 Solanum torvum fruits and leaf sample collection

Fresh fruits and leaves of *solanum torvum* were collected from the crop farms of the college of Agriculture Education (CAGRIC), University of Education, Winneba. The samples were conveyed to CAGRIC chemistry laboratory for extractions.

### 2.2 Extract preparation and administration

Aqueous extracts of *S. torvum* were prepared from the leaves and fruits of *Solanum torvum*. In the fruit sample, 100 grams (g) of *solanum torvum* fruit was blended (using an electric blender) with 100ml of distilled water, the mixture was sieved and the resulting filtrate obtained (150 ml) was labeled as STEFF (*Solanum torvum* extract fresh fruits). 100 g of the fresh leaves was also blended with 100 ml of distilled water, the resulting mixture was sieved and the filtrate obtained (150 ml) was labeled as STEFL (Solanum torvum extract fresh leaves). The extracts were administered to the respective experimental groups at 0.5 ml daily for a six weeks period while the extracts were refrigerated. New extracts were prepared every three days to avoid degradation in extract quality. The experimental animals were randomly divided into three groups with five replicates in each group. The animals were orally given 0.5ml daily for 6 weeks by syringe either of distilled water (Control), STEFF (Treatment 1) or STEFL (Treatment 2).

### 2.3 Experimental Animals and Feed

Fifteen (15) rabbits (aged 6-8 weeks) of both sexes and varying breeds were selected from two farmhouses in the Asante Mampong Municipality and were kept at the UEW Talif Project at the Animal farm of CAGRIC. The experimental animals were housed under standard conditions a week before the onset of the study and until the end. The animals were fed with standard concentrate and water *ad libitum* a weeks before the study. Concentrate (17.58 % crude protein) prepared from the mill house of the animal farm at UEW was used to feed the experimental animals during the study. The concentrate (100 kg) was formulated with; 75.0 kg wheat brand, 13.5 kg maize, 10.0 kg soya beans, 0.5 kg premix, 0.5 kg diacapsule sulphate and 0.5 kg salt.

### 2.4 Blood sample collection

1ml of Blood was taken from each experimental animal a week before the start of the experiment and after six weeks of treatment administration to access the effect of the treatments. All the blood samples were taken from the ears veins of the experimental animals using a 2ml Euro-ject-II syringe (Jiangyin N.M Product Ltd, China) and 25 G * 5/8 syringe (Neomedic PTY Ltd, South Africa). The blood samples were kept in EDTA tubes and frozen until further examination.

### 2.5 Determination of erythropoietic, hemanitic and leucocytic effects

The blood samples collected were conveyed to the hematology unit of Calvary hospital, Effiduase in the Ashanti region of Ghana. The total hematology components were determined using a BC 2600 Automatic Hematology Analyzer pre-centrifuged with a KMR-IV (Blood Mixer).

### 2.6 Statistical tool

Minitab (v. 17) and Microsoft excel were used to analyse the data by one-way ANOVA followed by Tukey’s comparison test and line graph respectively.

## 3. Results

### 3.1 Hemanitic effects of *Solanum torvum*

The hemoglobin levels of the rabbits in the control group and those treated with the aqueous extracts of raw fruits and leafs of *solanum torvum* were determined at the sixth week of treatment administration. The hemoglobin levels for all the treatments increased at week 6, which recorded a major rise in hemoglobin levels for animals treated with STEFF and STEFL with a clear difference between the control and the other treatments. Though the experimental animals of the control group were not anemic but the *S. torvum* treated groups should a significant increase in blood hemoglobin quality.

Grouping Information Using the Tukey Method and 95% Confidence

**Table 1.**

| Treatments |  | Hemoglobin (Hb) concentrations |  |
| --- | --- | --- | --- |
| Factor | N | Mean | Groupings |
| STEFL | 5 | 18.800 | A |
| STEFF | 5 | 16.68 | A |
| CONTROL | 5 | 12.400 | B |

One-way ANOVA (p<0.05) followed by Tukey’s comparison test. Means that do not share a similar letter are significantly different.

**Figure 1.**
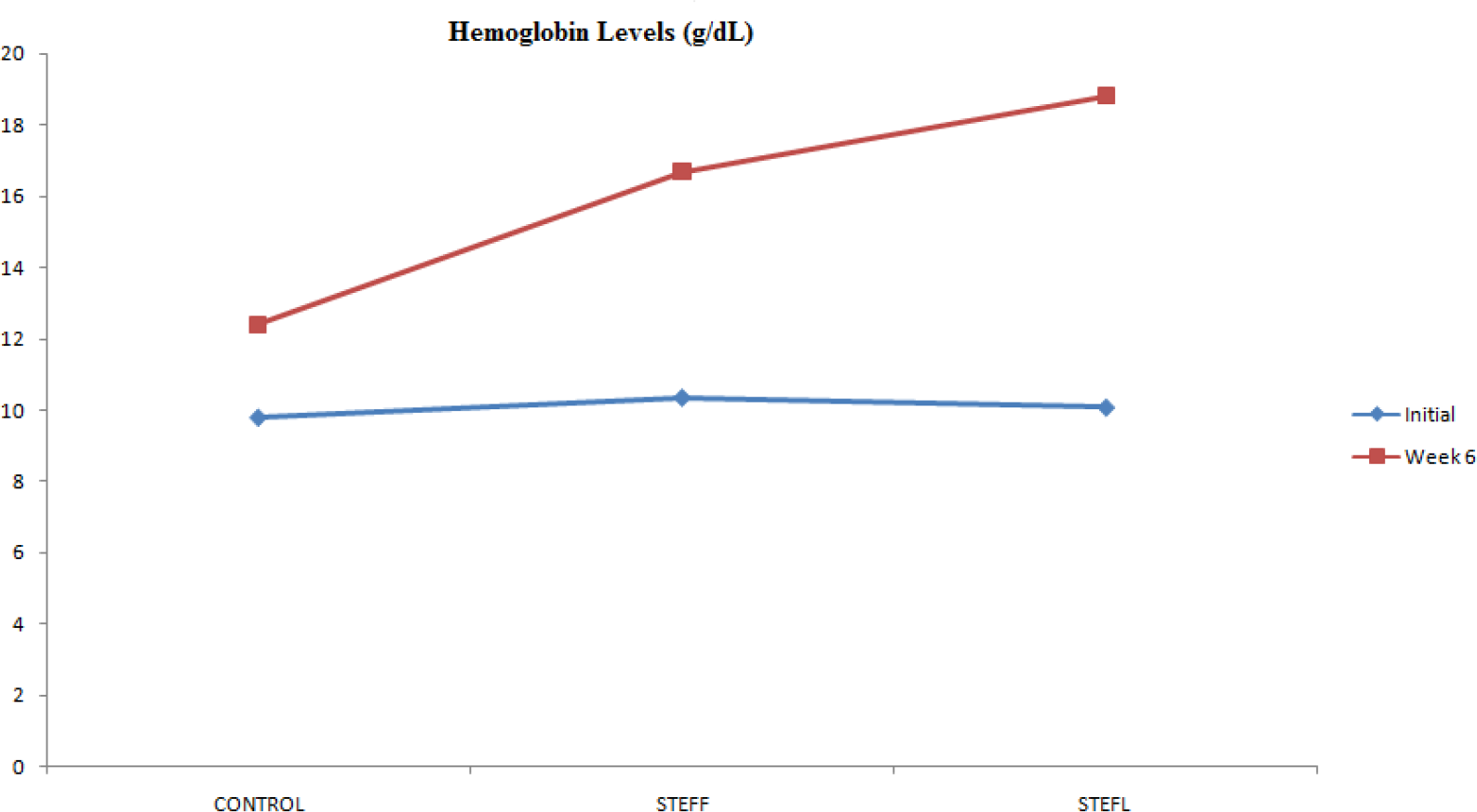
Blood hemoglobin level of experimental animals at initial and week six of treatment administration.

### 3.2 Effects of *Solanum torvum* on white blood cell counts

All groups demonstrated a rise in leucocyet count at the sixth week with the control recorging a marginal increase compared to the STEFL and STEFF administered groups. The white blood cell count of all the experimental animals were analysed at week six and analysed as follows;

**Table 2.**

| Treatment .1 | Mean WBC counts .2 |  |  |
| --- | --- | --- | --- |
| Factor | N | Mean | Grouping |
| STEFL | 5 | 9.550 | A |
| STEFF | 5 | 8.600 | A |
| CONTROL | 5 | 5.075 | B |

One-way ANOVA (p<0.05) followed by Tukey’s comparison test. Means that do not share a letter are significantly different.

**Figure 2.**
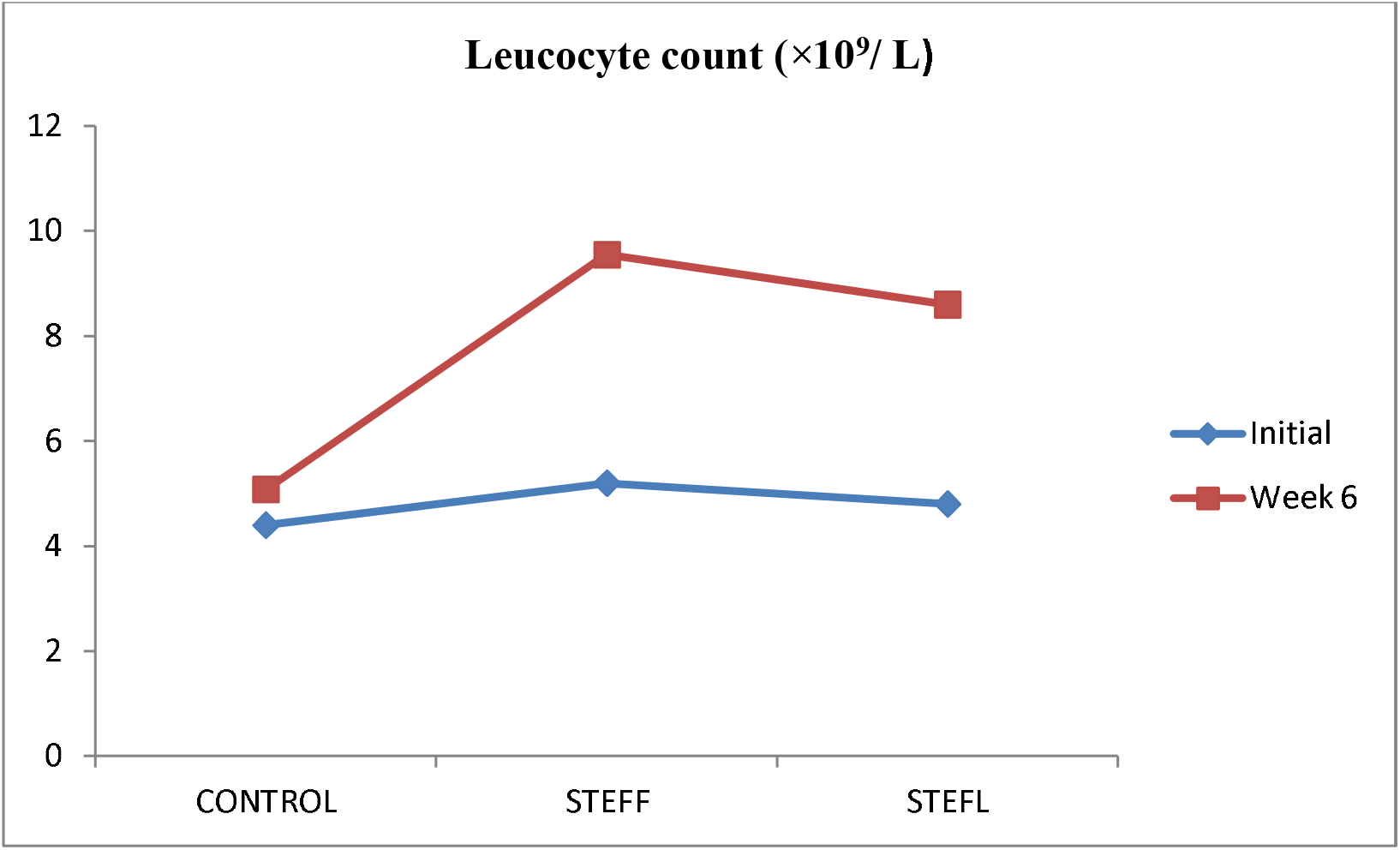
Leukocyte count experimental animals at initial and week six of treatment administration.

### 3.3 Erythropoietic effects of *Solanum torvum*

Increase in erythrocyte counts of the animals correspnded with increse in hemoglobin concentrations. The STEFL treated group recorded the highest rise in both hemoglobin and erythrocyte counts highlighting the correlation between the two parameters. The control also recorded a value which was within the normal range of erythrocyte count but was however below the counts of the two *S. torvum* treated groups. The result for eruthrocyte counts for the various groups are analysed below;

**Table 3.**

| Treatments | RBC |  |  |
| --- | --- | --- | --- |
| Factor | N | Mean | Grouping |
| STEFL | 5 | 6.630 | A |
| STEFF | 5 | 6.442 | A |
| CONTROL | 5 | 4.7500 | B |

One-way ANOVA (p<0.05) followed by Tukey’s comparison test. Means that do not share a letter are significantly different.

**Figure 3.**
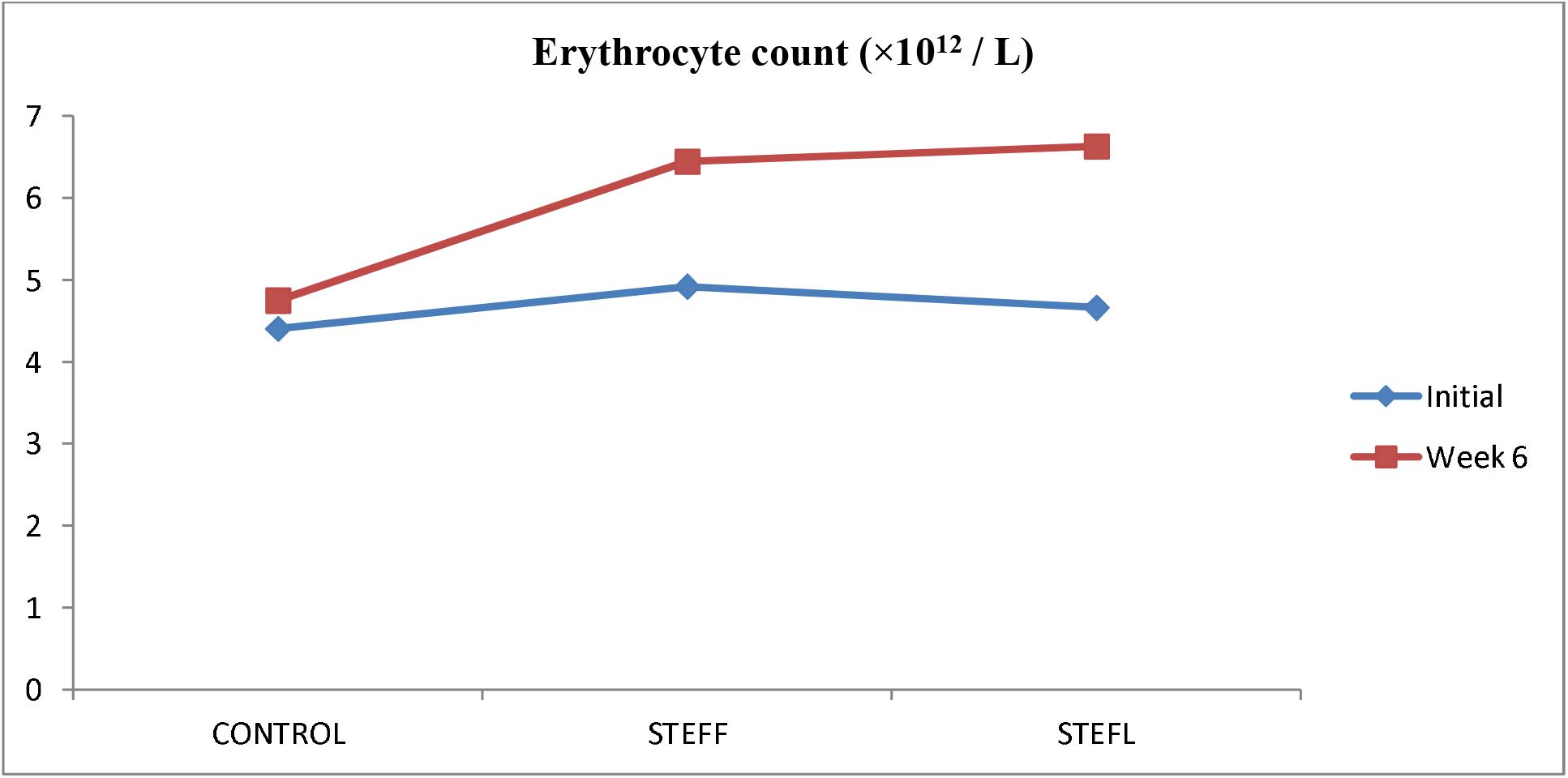
Erythrocyte count experimental of animals at initial and week six of treatment administration.

Increased RBC counts were recorded for all treatments in week 6. However, the animals treated with STEFF and STEBL recorded higher rise in erythrocyte counts relative to the control.

### 3.4 Effects of *Solanum torvum* on hematocrit

The hematocrit percentage for all treatments including the control increased at week 6. However, a significant difference was observed in the increments of the other treatments as compared to the control group. The STEFL treated group again recorded the highest rise in hematocrit which is in line with respective rises in hemoglobin concentrations and erythrocyte counts. The hematocrit represents the percentage of erythrocytes the blood and so any rise in erythrocyte count must correspond with rises in hematocrit.

The hematocrit percentage of the experimental animals are analysed as follows;

**Table 4.**
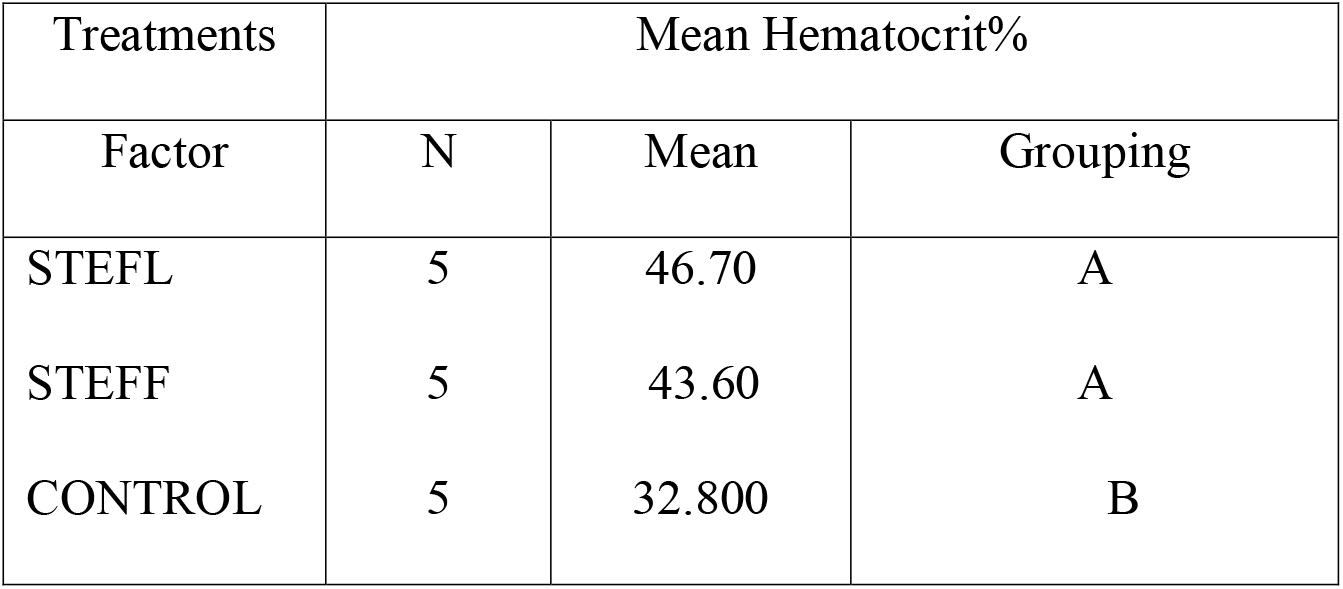

| Treatments | Mean Hematocrit% |  |  |
| --- | --- | --- | --- |
| Factor | N | Mean | Grouping |
| STEFL | 5 | 46.70 | A |
| STEFF | 5 | 43.60 | A |
| CONTROL | 5 | 32.800 | B |

One-way ANOVA (p<0.05) followed by Tukey’s comparison test. Means that do not share a letter are significantly different.

**Figure 4.**
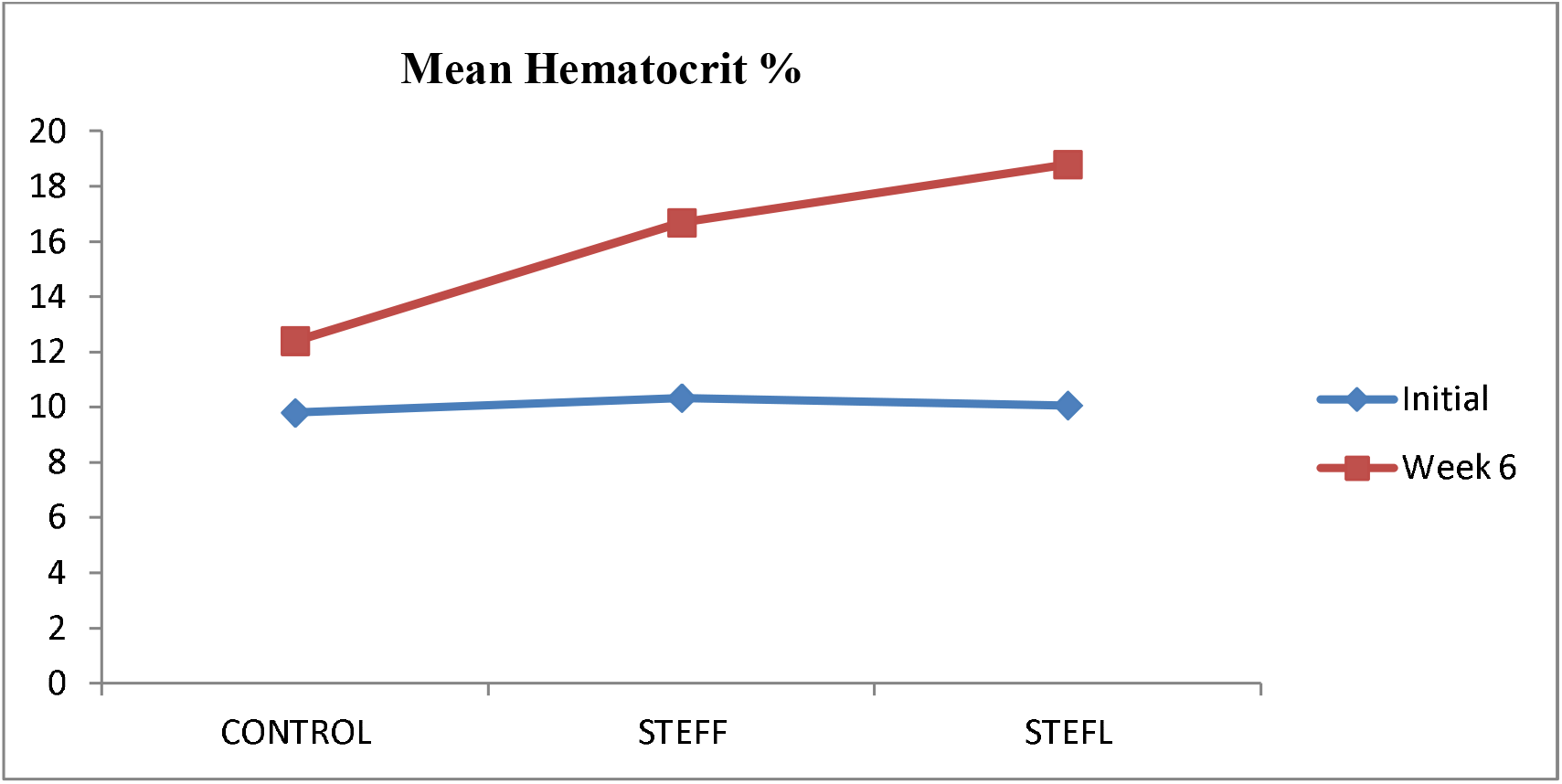
Blood hematocrit of experimental animals at initial and week six of treatment administration.

**Table 5.** Means and standard deviations of the experimental groups

| Treatments | WBC cells/mcL |  | HBG g/dL |  | RBC cells/mcL |  | HCT % |  |
| --- | --- | --- | --- | --- | --- | --- | --- | --- |
|  | mean | St.dv | mean | St.dv | mean | St.dv | mean | St.dv |
| Control | 5.1 | 1.1 | 12.4 | 0.3 | 4.8 | 0.1 | 32.8 | 1.7 |
| STEFF | 8.6 | 0.5 | 16.7 | 0.8 | 6.6 | 0.5 | 43.6 | 4.1 |
| STEFL | 9.6 | 0.9 | 18.8 | 2.1 | 6.4 | 0.7 | 46.7 | 3.3 |

## 4. Discussions

The hemoglobin (Hb) levels for all the treatments and the control increased at the sixth week and that confirms an increasing hemanitic activity for all treatments. However, all *S. torvum* treated group’s recorded higher Hb levels relative to the control group. Kuffuor *et al*. (2011) also demonstrated the erythropoietic effects of turkey berry plant in phenyl hydrazine induced anemic Sprague–Dawley rats. After six weeks of treatment administration, there was a significant recovery in blood hemoglobin levels. These hemanitic effects of *solanum torvum* can be associated with its richness in iron (76.869mg/kg), manganese (19.466 mg/kg), calcium (221.583 mg/kg), copper (2.642mg/kg), Akoto *et al*., (2015). Moreover, it contains other elements that are essential in the process of hematopoeisis. The rise in the hemoglobin levels of the control group is attributed to nutrients of the feed (concentrate) of the organisms however, the massive difference between the control and the other treatments are accounted for by *Solanum torvum* activities. The rise in mean WBC for all groups treated with the extract of *Solanum torvum* is accounted for by the effects of the extracts administered. Kuffuor *et al*., (2011) also demonstrated *Solanum torvum’s* Immunostimulatory activity and how it affected differential WBC count by immunizing Sprague–Dawley rats with Sheep Red Blood Cell before treating them with *Solanum torvum* extracts and found the extracts to cause significant increase in WBC counts relative to the non-treated groups. Chah *et al*., (2000) explained the antimicrobial and antifugal activity of *Solanum torvum* when they used its methanolic extracts against some bacteria species isolated from the clinical samples of humans. This activity of *Solanum torvum* can be explained by its leucocytic activity because the leucocytes are the major components involved in antigenic regulations. Kuffuor *et al*., (2011) also investigated this same effect by inducing anemia with PHZ in Sprague–Dawley rats and found *Solanum torvum* to cause significant increases in RBC count and hemoglobin levels after initially been decreased by 58.73% and 64.98% respectively. This observation also backed the results obtained in this study.

The general increase in hematocrit in the S. torvum treated groups is explained by the increases in RBC counts. The hematocrit percentage indicates the percentage of RBC in the total blood cells.

## 5. Conclusion

The data obtained makes it clear that *Solanum torvum* generally has an effect on hematopoeisis, hemanitic and leucocytic activities. However, *S. torvum* may have further effects on other blood components and properties that have not been discussed in this study.

